# Variable autoinhibition among deafness-associated variants of Diaphanous 1 (DIAPH1)

**DOI:** 10.1101/2021.03.10.434847

**Authors:** Rabina Lakha, Angela M. Montero, Tayyaba Jabeen, Christina C. Costeas, Jia Ma, Christina L. Vizcarra

## Abstract

One of the earliest mapped human deafness genes, *DIAPH1*, encodes the formin DIAPH1. To date, at least three distinct mutations in the C-terminal domains and two additional mutations in the N-terminal region are associated with autosomal dominant hearing loss. The underlying molecular mechanisms are not known, and the role of formins in the inner ear is not well understood. In this study we use biochemical assays to test the hypotheses that autoinhibition and/or actin assembly activities are disrupted by DFNA1 mutations. Our results indicate that C-terminal mutant forms of DIAPH1 are functional *in vitro* and promote actin filament assembly. Similarly, N-terminal mutants are well-folded and have quaternary structures and thermal stabilities similar to the WT protein. The strength of the autoinhibitory interactions varies widely among mutants, with the *ttaa*, A265S and I530S mutations having an affinity similar to WT and the 1213x and Δ*ag* mutations completely abolishing autoinhibition. These data indicate that, in some cases, hearing loss may be linked to reduced inhibition of actin assembly.

## Introduction

The sensory epithelia of the inner ear contain precisely-patterned cytoskeletal structures. These cytoskeletal networks include the paracrystalline array of actin filaments in mechanosensory stereocilia (Vélez-Ortega and Frolenkov, 2019), the actin and microtubule meshwork of the cuticular plate that anchors stereocilia (Pollock and McDermott Jr, 2015), and the tight junctions that maintain ionic composition of cochlear fluids (Wang *et al.*, 2015), among others. The importance of these cytoskeletal networks for auditory function is underscored by the large number of genes encoding cytoskeletal-associated proteins that are linked to hereditary deafness (Drummond *et al.*, 2012). These include genes encoding actin itself, as well as filament bundling proteins, nucleation factors, and myosin motors.

One of the first genes to be linked to inherited deafness was *DIAPH1* on chromosome 5 in humans. Variants of *DIAPH1* are associated with autosomal dominant, progressive, sensorineural hearing loss called DFNA1 (Lynch *et al.*, 1997). In the original report of DFNA1, the hearing loss is post-lingual, starting in the low frequency region and progressing to the entire frequency range (Lalwani *et al.*, 1998). The originally-described variant contains a single nucleotide change which creates a cryptic splice site. Four nucleotides, referred to below as ‘*ttaa*’, are inserted near the penultimate exon during mRNA processing (Lynch *et al.*, 1997), and the predicted protein sequence contains 21 spurious residues in place of the WT 52 residues (Figure 1A).

**Figure 1.**
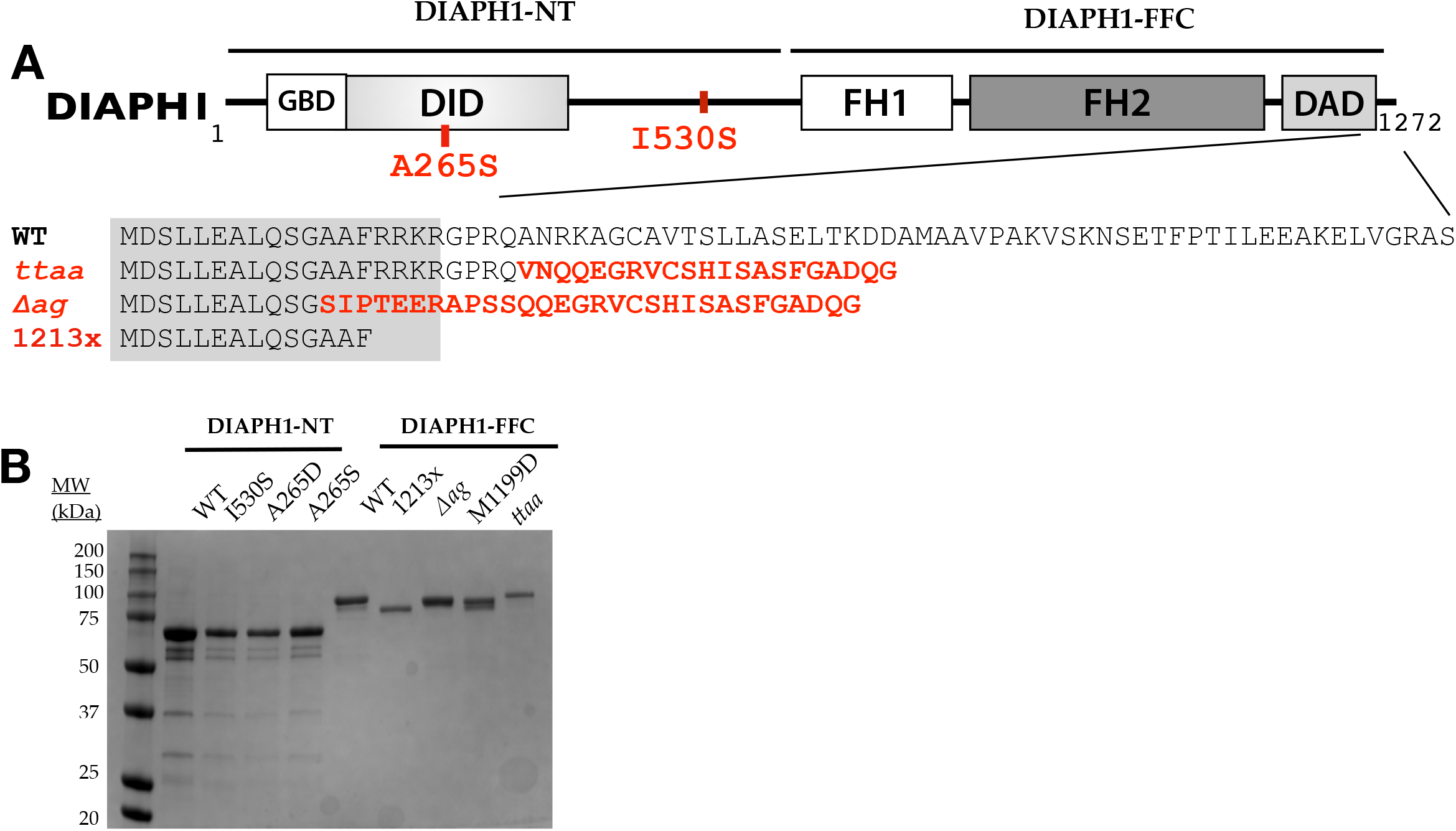
Isolation and structural analysis of DIAPH1 proteins. (A) Domain structure of the DIAPH1 protein, with DFNA1 mutations shown in red. Numbering is based on isoform NP_005210.3, with complete amino acid sequences provided in Table S1. The gray box over the sequence alignment shows the assigned boundary of the DAD domain. The ‘FFC’ region of the protein contains the FH domains and the C-terminus, and the ‘NT’ region encompasses the rest. Abbreviations: GBD, GTPase binding domain; DID, Diaphanous inhibitory domain; FH, formin homology; DAD, Diaphanous autoinhibitory domain. (B) SDS-PAGE analysis of purified proteins, shown with Coomassie staining.

The gene product, DIAPH1, is a member of the formin family of cytoskeletal regulators and, like other formins, interacts directly with the actin and microtubule cytoskeletons (Li and Higgs, 2003; Gaillard *et al.*, 2011). Formins contain highly conserved formin homology (FH) domains (Figure 1A). The FH2 domain binds actin monomers (Otomo *et al.*, 2010), actin dimers (Pring *et al.*, 2003), and actin filaments (Otomo *et al.*, 2005). This domain most likely also binds microtubules (Bartolini *et al.*, 2008) although less is known about the structure of formin/microtubule complexes. Members of the DIAPH sub-family of formins are regulated by Rho-GTPase binding (Lammers *et al.*, 2008). Rho binding to the DIAPH sub-family of formins disrupts an autoinhibitory intramolecular interaction between the N-terminal DID domain and the C-terminal DAD domain and enables these formins to then bind to actin (Lammers *et al.*, 2005; Li and Higgs, 2005; Wallar *et al.*, 2006). Among this sub-family, at least one other member, in addition to DIAPH1, has been associated with hearing loss: DIAPH3. A mutation in the 5’ untranslated region of *DIAPH3* is associated with an auditory neuropathy disorder, AUNA1 (Schoen *et al.*, 2010; Surel *et al.*, 2016).

Since the original linkage of *DIAPH1* to deafness in 1997, at least four additional distinct *DIAPH1* variants have been associated with autosomal dominant, progressive sensorineural hearing loss (Stritt *et al.*, 2016; Ueyama *et al.*, 2016; Ganaha *et al.*, 2017; Kang *et al.*, 2017, 2017; Neuhaus *et al.*, 2017; Kim *et al.*, 2019). The locations of these in the protein sequence are shown in Figure 1A. DFNA1 was originally classified as a non-syndromic form of deafness for the *ttaa* variant (Lalwani *et al.*, 1998), meaning that hearing loss was the only disorder. However, in recent studies, the R1213x and Δ*ag* (p.A1210S*fs*X31) variants of DIAPH1 were linked to autosomal dominant macrothrombocytopenia (MTP) in addition to hearing loss, suggesting that DFNA1 may be a syndromic form of deafness, at least for those mutations (Stritt *et al.*, 2016; Ganaha *et al.*, 2017; Neuhaus *et al.*, 2017; Westbury *et al.*, 2018). Inherited MTP is a disorder characterized by giant platelets with overall reduced circulating platelet counts (Noris *et al.*, 2014). Although individuals carrying the *ttaa* variant have normal lifespans (Lalwani *et al.*, 1998), it was suggested that blood smears be routinely analyzed for individuals with *DIAPH1* variants since individuals with R1213x and Δ*ag* variants were asymptomatic for MTP (Neuhaus *et al.*, 2017). It is unknown whether the R1213x knock-in mouse has platelet defects similar to MTP. Interestingly, it was recently reported that individuals with the A265S DIAPH1 variant have normal platelet size with severe to profound hearing loss (Kim *et al.*, 2019).

Three *DIAPH1* variants that lead to truncation prior to the FH2 domain have also been associated with autosomal recessive microcephaly, seizure disorders, and blindness (Ercan-Sencicek *et al.*, 2014; Al-Maawali *et al.*, 2015). These truncations are predicted to lead to non-sense-mediated mRNA decay, consistent with an observed lack of DIAPH1 expression in affected individuals (Ercan-Sencicek *et al.*, 2014). Complete knock-out of the mouse DIAPH1 homolog mDia1 results in myoproliferative defects similar to preleukemic state in humans (Peng *et al.*, 2007) and reduced T cell population in lymphoid tissues (Eisenmann *et al.*, 2007), but not microcephaly. Hearing thresholds were found to be normal in these mDia1 knockout mice (Ueyama *et al.*, 2016). Consistent with this, individuals with familial microcephaly linked to early truncation of the *DIAPH1* gene displayed no hearing loss (Ercan-Sencicek *et al.*, 2014; Al-Maawali *et al.*, 2015). These studies show that (1) DIAPH1 may not be essential for normal hearing in mouse or human, and (2) mouse mDia1 and human DIAPH1 may have divergent functions, complicating the use of mouse models for *DIAPH1*-linked diseases.

The native and pathological roles of formins in the auditory system are not well-understood. Due to their high-affinity, processive association with the barbed ends of actin filaments (Kovar and Pollard, 2004; Paul and Pollard, 2009; Courtemanche, 2018), formins are attractive candidates for localization to stereocilia tips. However, endogenous formin localization to the hair bundle has not been reported to date. In a knock-in mouse, expressing a fluorescently tagged DIAPH1-R1213x, localization of mutant DIAPH1 was observed in a subset of inner hair cell stereocilia tips (Ninoyu *et al.*, 2020). In the same study, endogenous Dia1 was localized most strikingly to apical junction complexes in inner hair cells, as well as in outer hair cells, Deiters’ cells, inner pillar cells, and spiral ganglion in both WT and transgenic mice at P5 (Ninoyu *et al.*, 2020). This study also revealed that WT mDia1 and mutant Dia1 (R1213X) were first observed in apical junction complexes of inner and outer hair cells at the basal turn which spread up to middle and apical turns as the Organ of Corti matures (from P5 to P14). In a separate study, immuno-localization of endogenous mDia1 in the inner ear was reported to be exclusively in inner pillar cells and the spiral ganglion in adult mice (Neuhaus *et al.*, 2017). Increased hair cell loss in 25-week transgenic mice expressing 1213x compared to WT mice suggests that DIAPH1 may be expressed in hair cells (Ueyama *et al.*, 2016). Given the wide-ranging localization data, it is currently challenging to match either WT DIAPH1 or variants of DIAPH1 to a specific cytoskeletal network within the cochlea.

Based on the dominant inheritance pattern and the location of the DFNA1 mutations in the DIAPH1 protein (Figure 1A), it has been proposed that the DFNA1 gene products are constitutively active and that this may be basis for their pathology. However, their variable position with respect to the known DID/DAD interface suggests that disruption of autoinhibition may be variable. Here we test this hypothesis for three C-terminal mutations and two N-terminal mutations using purified protein fragments and *in vitro* actin assembly assays. Our data show two classes of *DIAPH1* variants: those with undetectable auto-inhibitory interactions and those with auto-inhibition similar to WT DIAPH1. This data adds molecular context to the clinical heterogeneity of *DIAPH1*-associated hearing loss.

## Methods

### Protein expression constructs

All *DIAPH1* genes were constructed from clone BC117257 (Transomic Inc.). The translated sequence differs from protein NP_005210.3 by a nine amino acid deletion in the N-terminus, outside of the Rho-binding or DID domains, and a single proline deletion in the FH1 domain. We use the residue numbering of NP_005210.5 to be consistent with other reports in the literature. The complete amino acid sequences, including affinity purification tags, are given in Supplemental Information. Constructs of interest were cloned into pGEX6P2 vector for DIAPH1-NT, pMAL-c2x for INF2-NT, or the pET15b vector for FFC. A C-terminal strep tag was added to all FFC genes. Proteins were expressed in ROSETTA-DE3 *E. coli* cells (Novagen). For expression, transformed ROSETTA-DE3 cells were grown in 30– 60-mL Luria-Bertani Broth overnight at 37°C with shaking. Ten mL of this dense culture was used to inoculate 1 L of Terrific Broth. These large cultures were shaken at 250 rpm at 37°C until the optical density at 600 nm reached 0.3-0.6. The cells were then chilled to 18°C for at least 30 min, and protein expression was induced with 0.25 mM IPTG, followed by shaking overnight at 18°C. The cultures were harvested by centrifugation, resuspended in PBS (140 mM NaCl, 2.7 mM KCl, 10 mM Na_2_HPO_4_, 1.8 mM KH_2_PO_4_), flash frozen, and stored at –80°C. Unless specified below, all protein purification steps were carried out at 4°C, using ice-cold buffers.

### DIAPH1-NT purification

To isolate GST-tagged NT proteins, ≈25 g of frozen cells were thawed and brought to ≈35 mL with PBS supplemented with 1 mM PMSF, 4 μg/mL DNase (Sigma #DN25), and 1 mM DTT. The cells were lysed by French Press and the lysate was centrifuged with the speed of 30,000 – 40,000 ×*g*. The supernatant was incubated with 2 mL of glutathione sepharose resin slurry (Cytiva) for 1 h with rocking. The slurry was poured into a column, and the resin was washed with 25 mL of PBS. The protein was then eluted from the column with glutathione elution buffer (50 mM Tris-Cl, 20 mM glutathione, 50 mM NaCl, pH 8) and dialyzed in PBS for 3 h. GST was cleaved from the NT protein using PreScission Protease by incubating overnight in dialysis tubing. To remove GST, the sample was nutated with glutathione sepharose, and the unbound fraction was collected. The cleaved protein was dialyzed twice in 1 L MonoQ A buffer (10 mM Tris, 1 mM DTT, pH 8.0) for 1.5 h each. The protein was filtered through a 0.2 μm PES filter and loaded onto a MonoQ 5/50 column (Cytiva). Then, the column was washed with 10 column volumes (CV) of 25% MonoQ B buffer (10 mM Tris, 1 M KCl, 1 mM DTT, pH 8.0) followed by a step elution from 25-65% MonoQ B buffer. The peak fractions were pooled based on SDS-PAGE analysis. The pooled protein fractions were buffer exchanged via PD-10 desalting column (Cytiva) into storage buffer (10 mM Tris, 100 mM KCl, 1 mM DTT, pH 8.0), flash frozen in liquid nitrogen, and stored at –80°C until used in kinetic and other assays.

### INF2-NT purification

INF2-NT cells were lysed in Extraction buffer (200 mM NaCl, 20 mM, sodium phosphate, 1 mM EDTA, 1 mM DTT, pH 8.0) in French Press and the supernatant was collected after centrifugation. The supernatant was rocked with 2 mL of amylose resin slurry (NEB) for 1 h. The protein was then eluted in 200 mM NaCl, 20 mM sodium phosphate, 10 mM maltose, 1 mM EDTA, 1 mM DTT, pH 8.0. The protein was then cleaved overnight with 30 μL of PreScission Protease by incubating overnight in dialysis tubing in the extraction buffer. The unbound protein was then nutated with amylose resin for 1 h and the unbound fraction was collected using a gravity column. The unbound protein was dialyzed twice in 1 L MonoQ A buffer for 1.5 h each. The protein was filtered through a 0.2 μm PES filter, loaded onto a MonoQ 5/50 column, and eluted with a linear gradient of 0-100% MonoQ B buffer over 30 CV. The pooled fractions were loaded onto a Q-HP anion exchange column (Cytiva) and eluted with a step to 40% MonoQ B buffer. The purified INF2 protein was aliquoted in small fractions and flash frozen in liquid nitrogen before storing at –80°C. Concentration was calculated using the absorbance at 280 nm and a predicted native extinction coefficient of 26,930 M^−1^cm^−1^.

### DIAPH1-FFC purification

To isolate all FFC proteins except 1213x-FFC, cells were lysed using french press in Ni-NTA lysis buffer (500 mM NaCl, 50 mM NaPi, 1 mM EDTA, 1 mM DTT, 1 mM PMSF and 4 μg/mL DNase pH 7.5). The supernatant was collected after centrifugation of the lysate and nutated with ≈800 μL of NI-NTA resin slurry (Thermo Scientific) for 45 min. The mixture was then passed through the gravity column and washed with Ni-NTA lysis buffer followed by washing with 500 mM NaCl, 50 mM NaPi, 10 mM imidazole, 1 mM EDTA, 1 mM DTT, pH 7.5. The protein was eluted with 500 mM NaCl, 50 mM NaPi, 400 mM imidazole, 1 mM EDTA and 1 mM DTT, pH 7.5. Elution fractions were buffer exchanged into strep wash buffer (150 mM NaCl, 100 mM Tris-Cl, 1 mM EDTA, 1 mM DTT, pH 7.5) in a PD-10 column before rocking with 500μL of Strep-T actin Sepharose resin slurry (IBA Lifesciences) for 45 min. After nutating the mixture, the protein was eluted using strep elution buffer (150 mM NaCl, 100 mM Tris-Cl, 2.5 mM Desthiobiotin, 1 mM EDTA, 1 mM DTT, pH 7.5).

For purifying 1213x-FFC, cells were resuspended in TALON extraction buffer (300 mM NaCl, 50 mM NaPi, 1 mM β-mercaptoethanol (bME), pH 8.0) supplemented with 1 mM PMSF and 4 μg/mL DNase. After lysis and centrifugation as described above, the supernatant was nutated with 2 mL of TALON resin (Takara) for 45 min. The resin was washed with TALON wash buffer (300 mM NaCl, 50 mM NaPi, 1 mM bME) and eluted with TALON elution buffer (300 mM NaCl, 50 mM NaPi, 200 mM Immidazole,1 mM bME, pH 7.0). The eluted protein was buffer exchanged with MonoQ A buffer using a PD-10 column, loaded onto a MonoQ 5/50 column, and eluted from the ion exchange column with a gradient of 0 to 400 mM KCl over 30 CV.

For all FFC proteins, following the final purification column, the protein was buffer exchanged using a PD-10 desalting column equilibrated with PBS (supplemented with 1 mM DTT). Protein aliquots were frozen in liquid nitrogen and stored at –80°C until assays were performed. For DIAPH1-NT and DIAPH1-FFC proteins, concentrations were calculated using SDS-PAGE and densitometry with the quantitative protein stain SYPRO Red (Invitrogen).

### Actin purification and labeling

To isolate actin, 3 g of rabbit muscle acetone powder (Pel Freez Biologicals) was mixed with 100 mL of actin extraction buffer (2 mM Tris-HCl, 0.2 mM CaCl_2_ 0.2 mM ATP,1 mM DTT) and stirred for 30 mins. The mixture was then filtered using a cheese cloth and the filtrate was collected. The extraction process was repeated on the crude residue. The combined filtrates were centrifuged for 1 h at 40,000 ×*g*. The supernatant was syringe filtered through a 0.2 μm PES membrane. The filtered actin solution was polymerized by stirring for 1.5 h with 50 mM KCl, 2 mM MgCl_2_, and 1 mM Mg-ATP. The KCl concentration was brought to 600 mM and stirred for an additional 30 min. The polymerized actin was then centrifuged for 1 h at 170,000 ×*g*. The supernatants were decanted and the pellets were soaked overnight with 0.5 mL actin extraction buffer. The soaked pellets were broken up with a dounce homogenizer. The actin was then dialyzed for 3-5 days in G-buffer (2 mM Tris-Cl, pH 8.0, 0.1 mM CaCl_2_, 0.2 mM ATP, 0.5 mM TCEP, 0.04% sodium azide). The dialyzed sample was centrifuged for 1-2 h at 170,000 ×*g* at 4°C and injected onto a HiLoad 16/600 Superdex 75 pg column (Cytiva) equilibrated with G-buffer.

For pyrene labeling, actin was polymerized for 30 min at 25°C in KMEH (10 mM HEPES, pH 7.0, 1 mM EGTA, 50 mM KCl, 1 mM MgCl_2_). The F-actin was then dialyzed (2 × 1.5 h, 4°C) into 0.1 M KCl, 25 mM imidazole, 2 mM MgCl_2_, 0.1 M ATP, pH 7.5 in order to remove reducing agents. A DMSO solution with ≈4–7 mol pyrene-iodoacetamide (Molecular Probes) per mol of actin was added to the F-actin mix. The reaction was stirred overnight and quenched by adding 10 mM DTT to the F-actin. The labeled F-actin was centrifuged at 180,000 ×*g* for 2 h. The pellets were resuspended in 4 mL G-buffer and the pellets were broken up with a dounce homogenizer. The actin was dialyzed for 2–3 days in G-buffer. The dialyzed sample was then centrifuged at 170,000 ×*g* for 2 h and injected onto a HiLoad 16/600 Superdex 75 pg column equilibrated with G-buffer. Unlabeled actin concentration was determined by measuring native absorbance at 290 nm (∊290nm = 26,596 M^−1^cm^−1^). For pyrene-labeled actin, pyrene concentration was calculated using the extinction coefficient 21,978 M^−1^cm^−1^, and the actin concentration was calculated using the correction factor of 0.127 at 290 nm ([actin] = (A_290 nm_ – 0.127× A_344 nm_)× 38 μM).

### Profilin purification. Schizosaccharomyces pombe

profilin was expressed in BL21-DE3 cells. Frozen cells were resuspended in lysis buffer (10 mM Tris-Cl, 1 mM EDTA, 1 mM DTT, 1 mM PMSF, pH 7.5), lysed by French Press, and centrifuged at 95,000 ×*g* for 20 min. The clarified cell lysate was passed over DE52 anion exchange resin and washed with lysis buffer. The unbound fraction was collected. Protein was first precipitated by bringing the ammonium sulfate concentration to 35% saturation and stirring at 4°C for 20 min. The precipitate was cleared by centrifugation at 25,000 ×*g* for 30 min. The profilin was precipitated by bringing the ammonium sulfate concentration to 61% and stirring at 4°C for 20 min, following centrifugation at 30,000 × *g* for 30 min. The pellet containing profilin was resuspended in 20 mL dialysis buffer (5 mM potassium phosphate, 1mM DTT, pH 7.5) and dialyzed overnight. The sample was filtered through a 0.2 μm PES membrane, and passed over a Mini CHT hydroxyapatite Type 1 column (Bio-rad). The unbound fraction was collected and concentrated using a Vivaspin 6 (3 kDa MWCO; Cytiva) to a final volume of 10 mL. The concentrated protein was filtered through a 0.2 μm PES membrane and gel filtered on a HiLoad 16/600 Superdex 75 pg column, equilibrated with 10 mM HEPES, 150 mM NaCl, 0.5 mM TCEP, pH7. The collected fractions were then pooled together, concentrated using an Amicon Ultra unit (3000 MWCO; Merck Millipore Ltd.) at speed of 3500 ×*g* to bring the final volume to 2 mL. The concentration of profilin was determined by measuring absorbance at 280 nm with an extinction coefficient of 21,860 M^−1^cm^−1^ (Lu and Pollard, 2001).

### Biophysical characterization

Circular dichroism (CD) spectra of DIAPH1-NT proteins were measured using Chirascan V100 Circular Dichroism Spectrometer from Applied Photophysics. Experiments were performed at room temperature in 1× PBS, in a quartz cuvette (1 mm path length). Spectra in Figure 3 are an average of three scans from 190 to 300 nm wavelength with a 1 nm step. For thermal denature experiments, the CD signal (*θ*) at 222 nm was recorded for every temperature from 10° to 70° with step of 1°C at the rate of 0.5 °C /min and 0.2 tolerance. The molar ellipticity was calculated based on the concentrations of proteins using [θ] = 100×*θ*/(*C×l*) where *C* is the molar concentration, and *l* is the cell pathlength in cm.

Size Exclusion with Multi Angle Light Scattering (SEC-MALS) was run on an Agilent 1260 Infinity II HPLC system, including a DAD detector at wavelength 280 nm, coupled with a Wyatt DAWN detector and an Optilab detector. A 100 μL sample was loaded onto a Superdex 200 10/300 column with 1× PBS (pH 7.4) as the running buffer at a 0.5 mL/min flow rate. The ASTRA software V7.3.0.18 was used to collect data and analyze results.

### Actin assembly assays

Bulk actin assembly assays were carried out essentially as described (Zuchero, 2007). Rabbit skeletal muscle was stored in G-buffer (2 mM Tris-Cl, pH 8.0, 0.1 mM CaCl_2_, 0.2 mM ATP, 0.5 mM TCEP, 0.04% sodium azide). Prior to polymerization, a mix of actin with 5% pyrene-labeling was incubated for 2 min at 25°C with 200 μM EGTA and 50 μM MgCl_2_ to convert Ca-G-actin to Mg-G-actin. When included in the experiment, profilin (typically 2:1 profilin:actin) was added to actin before conversion to Mg-G-actin. Polymerization was initiated by adding polymerization buffer (50 mM KCl, 1 mM MgCl_2_, 1 mM EGTA, 10 mM HEPES pH 7) to the Mg-actin. Pyrene fluorescence was measured every 15 s using a TECAN F200 plate reader with excitation filter at 360 nm (35 nm bandwidth) and an emission filter of 415 nm (20 nm bandwidth).

## Results

### WT and mutant DIAPH1-FFC accelerate actin assembly

At least three C-terminal variants and two N-terminal variants of DIAPH1 are associated with autosomal dominant progressive hearing loss (Figure 1A). To study autoinhibition in these variants, mutant and WT DIAPH1-FFC and NT proteins were recombinantly expressed and affinity purified (Figure 1B). FFC proteins, which include the FH1 and FH2 domains and the C-terminal region, were purified using a double-tagged purification to prevent degradation at the C-terminus that could confound the comparison of mutants (described in Methods). Using the pyrene-actin polymerization assay (Cooper *et al.*, 1983), we first tested the ability of WT and mutant FFCs to stimulate actin filament assembly in the presence and absence of the actin monomer binding protein profilin (Figure 2). All FFC proteins stimulated filament assembly to a degree similar to what has been observed for the mouse homolog mDia1 (Li and Higgs, 2003; Gould *et al.*, 2011): low nanomolar concentrations of FFC induced rapid polymerization, allowing the filamentous actin to reach a steady concentration within approximately 600 s.

**Figure 2.**
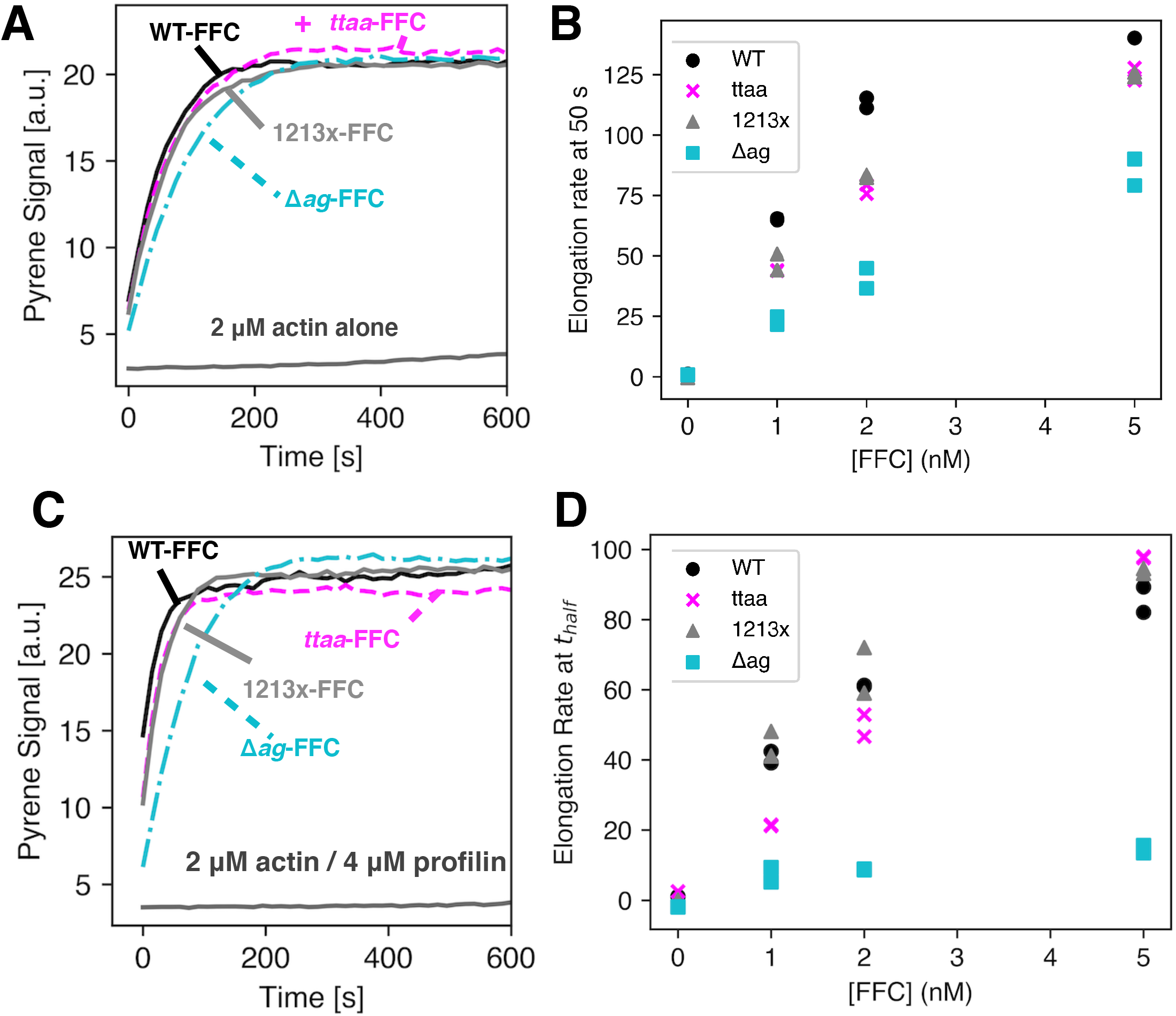
Actin assembly assays with DIAPH1-FFC. Pyrene-actin assembly assays to compare the nucleation activity of 5 nM WT or mutant FFCs with (A) 2 μM actin or (C) 2 μM actin and 4 μM *S. pombe* profilin. (B) Elongation rate at 50 seconds for pyrene-actin assay with 2 μM actin alone at 0, 1, 2 and 5 nM concentrations of FFC. (D) Elongation rate at time to half maximal polymerization (t*half*) for 2 μM actin and 4 μM *S. pombe* profilin at 0, 1, 2 and 5 nM concentrations of FFC. In all assays, 5% of the actin monomers were pyrene-labeled.

Compared to WT-FFC, the Δ*ag* mutation showed the only notable decrease in activity (Figure 2). Because assembly was so much slower in the presence of profilin, the slope at the time to half maximal polymerization (t_half_) was used to compare polymerization rates (Figure 2D). Of the FFC mutations, the Δ*ag* mutation leads to the removal of more WT residues in the DAD (Figure 1A). It has been observed that truncation of the entire DAD of mDia1 decreases actin nucleation activity (Gould *et al.*, 2011) although the specific residues that are disrupted in the Δ*ag* mutation have not been studied previously. Because this specific frameshift mutation also leads to 30 additional residues in the translated sequence, we cannot rule out interference of these residues in the formin/actin interaction. Alternatively, this more severe disruption may destabilize the FFC proteins. Indeed, all purified FFC proteins were found to be stable *in vitro* at –80°C for several weeks only.

### N-terminal mutants have biophysical properties similar to WT DIAPH1-NT

Two deafness-associated mutants in the N-terminus of DIAPH1 have been recently discovered: A265S (Kim *et al.*, 2019) and I530S (Kang *et al.*, 2017). The A265S mutant is located in the interface between the DID and DAD domains. The analogous residue position in mouse Dia1 (Ala256) was mutated to Asp based on the x-ray crystal structure of the DID/DAD complex, completely disrupting DID/DAD binding as measured by isothermal titration calorimetry (Lammers *et al.*, 2005). The other N-terminal DFNA1 mutant, I530S, is located in the predicted coiled coil domain (Figure 3A). Specifically, Ile530 is a hydrophobic amino acid in the heptad repeat highlighted in Figure 3A (Kang *et al.*, 2017), indicating that it could stabilize the coiled coil dimer. Given the putative structural role of isoleucine 530, we wanted to assess the stability and quaternary structure of I530S DIAPH1-NT in comparison to WT and A265S or A265D mutants.

**Figure 3.**
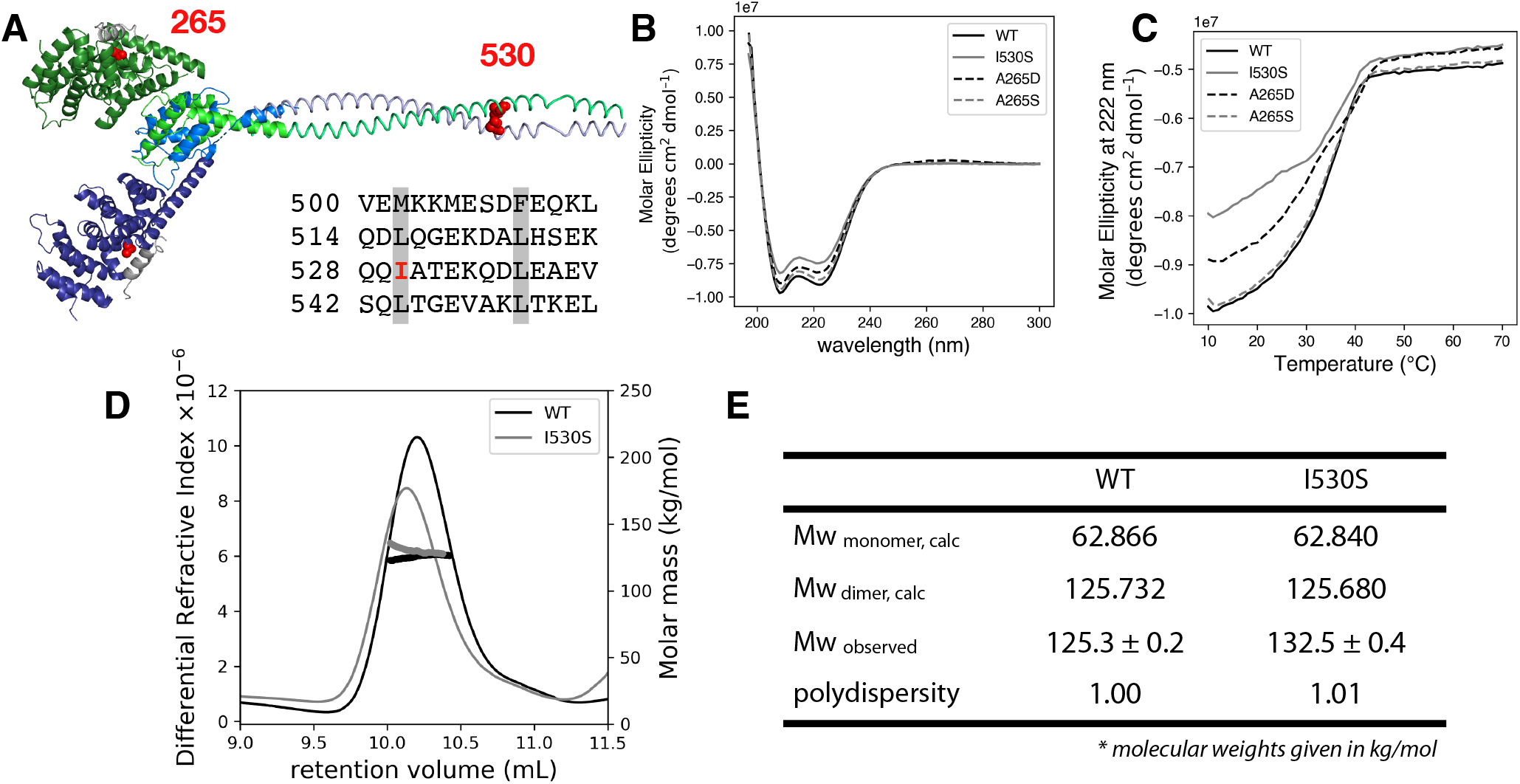
Structural characterization of NT mutants. (A) The predicted DID and dimerization domain structure of DIAPH1 were constructed from the PDB files 2BNX and 2F31 and from the prediction program COILS (Lupas *et al.*, 1991). The DD sequence is shown with the heptad repeat of hydrophobic residues highlighted in gray. The two DID-DD chains are each displayed in shades of blue and green. The DAD helices are shown in gray, and mutated residues 265 and 530 are in red spheres. Circular dichroism (B) wavelength scan and (C) thermal denaturation monitored at 222 nm for DIAPH1-NT WT and mutants. Conditions: 3 μM protein in phosphate-buffered saline. SEC-MALS analysis was done on both WT and the I530S mutant of DIAPH1. (D) Plot of differential refractive index and molecular mass (thick line) vs elution volume. The observed average molecular weights in panel (E) are closest to the dimer calculated Mw. The SEC-MALS data is representative of two trials, with separate protein preparations. In the other trial, Mw/polydispersity: WT, 133.7/1.04; I530S, 129.6/1.18.

All four versions of DIAPH1-NT were recombinantly expressed, affinity purified as GST fusion proteins, and then cleaved from their GST tags (Figure 1B). The four versions include WT, the two deafness-associated variants (A265S and I530S), and one engineered mutation (A265D, described above). The WT and three mutated proteins showed high alpha-helical content in their CD spectra, with minima around 208 and 222 nm (Figure 3B). Thermal denaturation was monitored at 222 nm, and the curves had inflection points between 38°C and 40°C (Figure 3C). For all variants, thermal unfolding was irreversible. Because the I530S mutation lies at a potentially important site for dimer stability, we analyzed this variant with SEC-MALS, which provides a shape-independent measurement of protein mass in solution (Minton, 2016). For both WT and I530S DIAPH1-NT, the major species in solution was closest to the calculated dimer molecular weight (Mw) of ≈125 kDa. No peak was observed near the predicted monomer Mw. While we cannot rule out the possibility the I530S undergoes rapid exchange with the monomer in solution, our data supports the assertion that all variants and the WT are dimers in solution and that the DIAPH1-NT variants studied have biophysical properties similar to the WT protein.

### Autoinhibition is variable among DFNA1 variants

The autosomal dominant inheritance pattern of DFNA1-associated deafness supports a gain of function for DIAPH1 variants. The location of the mutations, along with the induction of the extended microvilli in HeLa cells expressing DIAPH1-1213x (Ueyama *et al.*, 2016), points to a loss of the auto-inhibitory interaction between the DID and DAD domains. To test this, we titrated WT DIAPH1-NT into the pyrene-actin assays with different DIAPH1-FFC variants (Figures 4 and S2). These experiments were done under conditions favoring actin filament nucleation (2 μM actin, no profilin). The 1213x and Δ*ag* variants showed a complete loss of auto-inhibitory interactions (Figure 4C, D) similar to the M1199D variant (Figure 4E). The M1199D mutation is analogous to the M1182D mutation in mDia1, that has been previously shown to completely disrupt the DID/DAD (Lammers *et al.*, 2005). In contrast, the *ttaa* mutation was inhibited by DIAPH1-NT (Figure 4B). An overlay of all titrations plotted as % activity (Figure 4F), shows that the mutants cluster into two groups: 1213x and Δ*ag* form a constitutively activated group, while the actin filament nucleation activity of *ttaa* FFC can be inhibited by DIAPH1-NT with an inhibition constant that was indistinguishable from the WT. Because intermolecular DID/DAD interactions have been reported between mDia1 and the formin INF2 (Sun *et al.*, 2011), we also tested inhibition by the N-terminal fragment of INF2. We found a similar trend for inhibition of DIAPH1 FFC variants with INF2-NT (Figure S1) and DIAPH1-NT (Figure 4): complete auto-activation of 1213x and Δ*ag* and robust inhibition of WT and *ttaa*.

**Figure 4.**
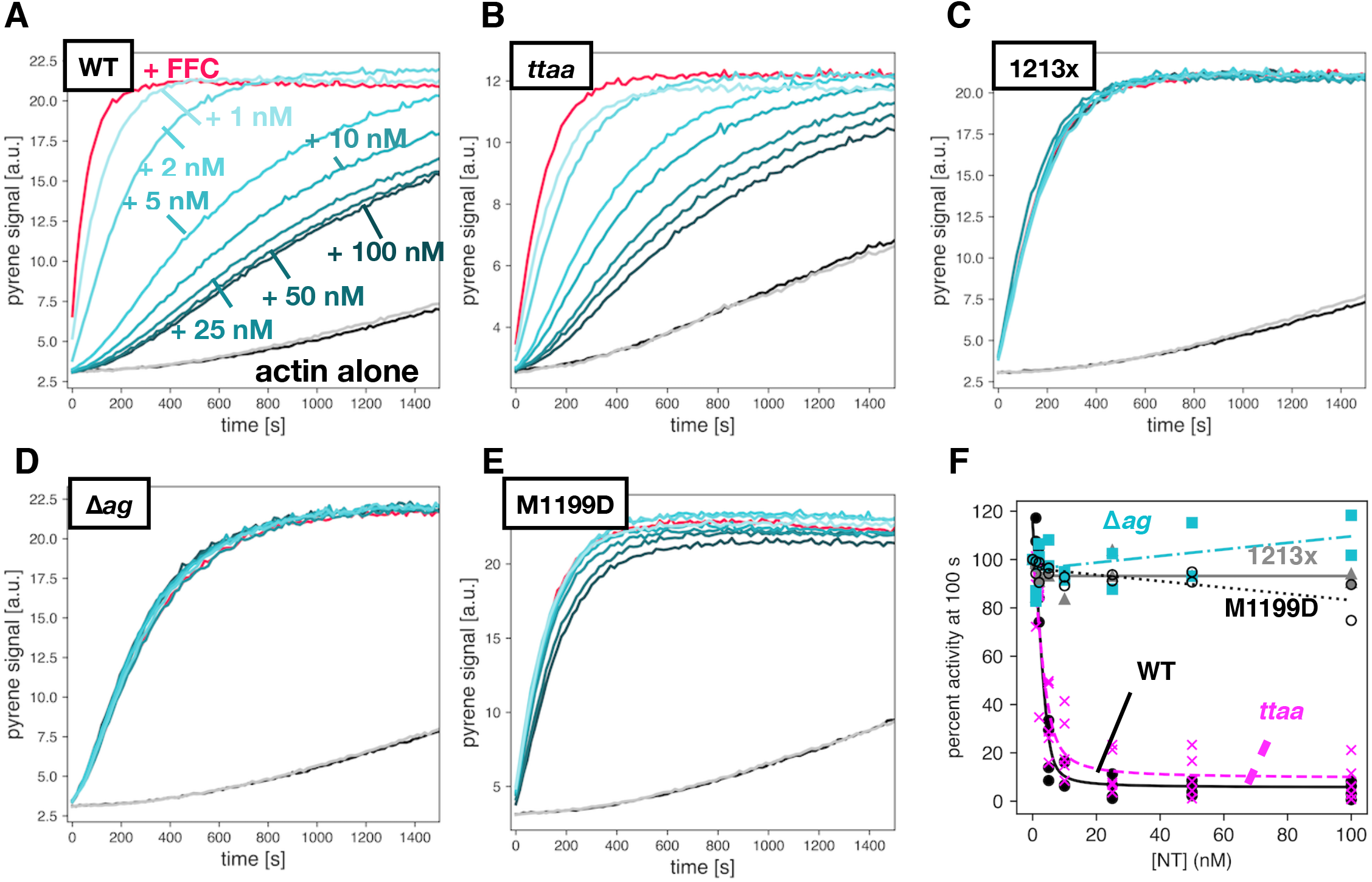
Varied auto inhibition of DIAPH1-FFC mutants. Pyrene-actin assembly assays were carried out with 2 μM actin (5% pyrene-labeled), ± 5 nM of FFC, either (A) WT; (B) *ttaa*;(C) 1213x; (D) Δ*ag*; or (E) M1199D, with varied concentration of WT DIAPH1-NT. Concentrations of NT are indicated in (A), and the color scheme is the same in panels (B-E), with actin plus 200 nM NT and no FFC in gray. (F) Inhibition curves calculated from raw pyrene traces and fit with a quadratic binding model that assumes 1:1 binding between dimers. A replicate of this experiment with completely different protein preparations is shown in Figure S2.

We next measured the inhibitory activity of DIAPH1-NT variants against WT-FFC. Using A265D as a no interaction control (Lammers *et al.*, 2005), we found that both A265S and I530S retained some autoinhibitory interaction (Figures 5 and S3). In all of our experiments, when titrating DIAPH1-NT into FFC, we observed residual FFC activity even at very high concentration of NT with inhibition plateaued at ≈10% of the elongation rate of FFC without NT, instead of ≈1-2% as observed previously with mouse isoforms (Li and Higgs, 2005). It is not clear whether this is a meaningful difference of between the mouse and human isoform. The residual activity of the FFC domain at high [NT] is particularly apparent in the A265S titration curve (Figure 5B) and could represent a DID-bound conformation where additional actin monomers can associate with the FH2 domain (Otomo *et al.*, 2010).

**Figure 5.**
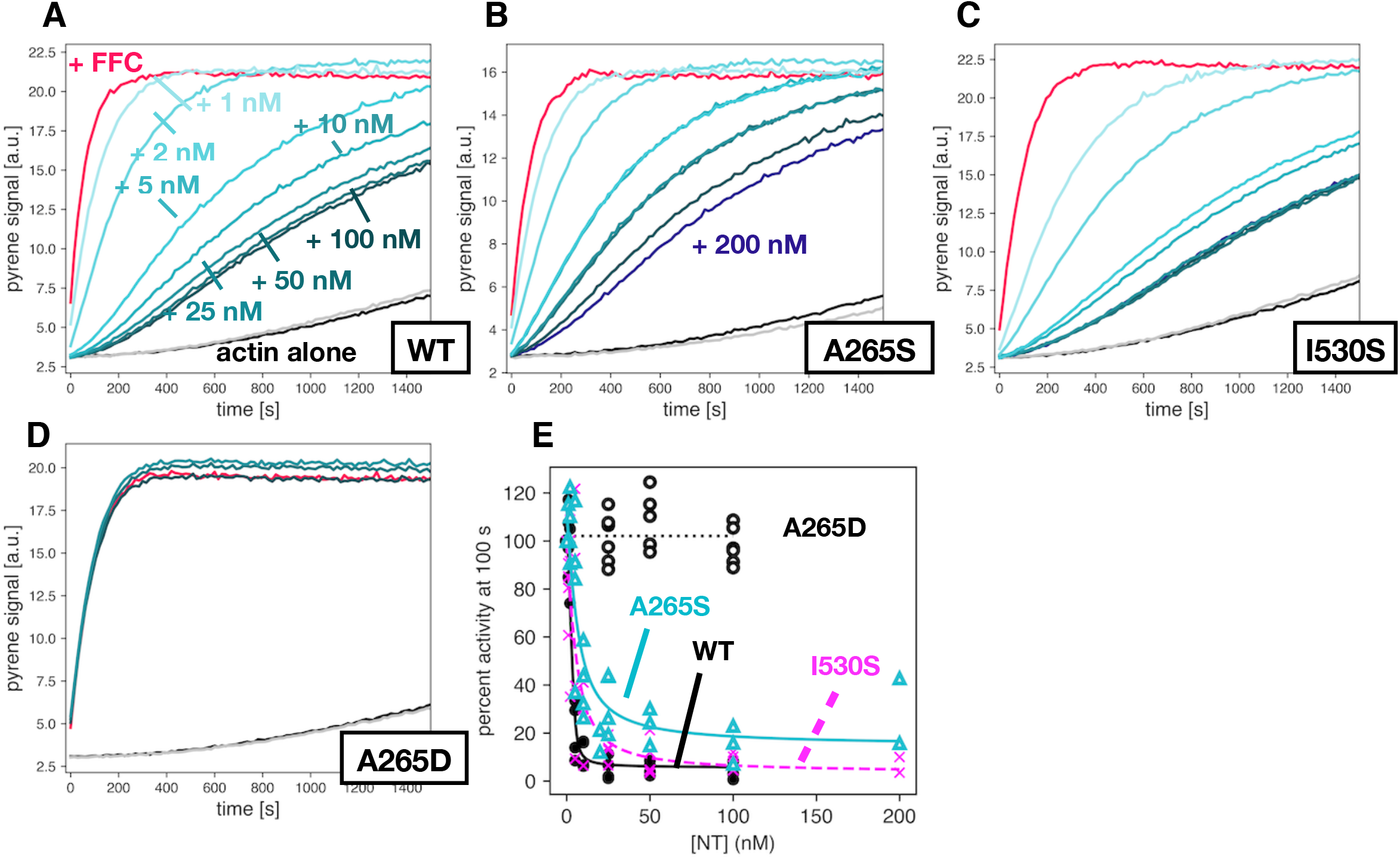
Auto inhibition by DIAPH1-NT mutants. Pyrene-actin assembly assays were carried out with 2 μM actin (5% pyrene-labeled), ± 5 nM of WT-FFC, titrated with either (A) WT; (B) A265S; (C) I530S; or (D) A265D DIAPH1-NT. Concentrations of NT are indicated in (A), and the color scheme is the same in panels (B-E), with actin plus 200 nM NT and no FFC in gray. (F) Inhibition curves calculated from raw pyrene traces and fit with a quadratic binding model that assumes 1:1 binding between dimers. Note that panel (A) in Figures 4 & 5 is the same plot and is shown twice to facilitate comparison with mutants. A replicate of this experiment with completely different protein preparations is shown in Figure S3.

## Discussion

Our data show that some *DIAPH1* variants associated with DFNA1 hearing loss retain the interactions that lead to autoinhibition. Based on this biochemical analysis, DFNA1 mutations can be grouped into at least two categories: (1) the *ttaa*, A265S and I530S variants, which retain measurable autoinhibitory interactions and (2) the 1213x and Δ*ag* variants, which completely abolish autoinhibition. Our data is consistent with the known importance of the basic ‘RRKR’ motif in the DID/DAD binding (Wallar *et al.*, 2006).

This classification is also consistent with a recent study from Kim *et al.*, in which the A265S mutation was mapped in a Korean family with non-syndromic, autosomal dominant, sensorineural hearing loss (Kim *et al.*, 2019). Using *in vitro* pull-down experiments and localization in HeLa cells, they build a model in which A265S could have “milder pathogenic potential” than 1213x, as A265S should showed intermediate DID/DAD association and mis-localization, compared to WT and 1213x (Kim *et al.*, 2019). They speculate that this could be correlated to the normal blood smears in individuals with the A265S variant: milder pathogenicity is associated with non-syndromic deafness, while 1213x is associated with both deafness and macrothrombocytopenia. It is notable that blood smears have not been reported for individuals with the I530S and *ttaa* variants. If these variants are non-syndromic like A265S, then given reports of MTP in individuals with 1213x and Δ*ag* variants (Stritt *et al.*, 2016; Neuhaus *et al.*, 2017; Westbury *et al.*, 2018), the classifications of syndromic versus non-syndromic DFNA1 would coincide with the two groups described in this report. Furthermore, Ueyama *et al.* observed minimal actin stress fiber formation in HeLa cells transfected with *ttaa* variant, while 1213x induced stress fibers to a degree similar to the M1199D mutant (Ueyama *et al.*, 2016).

In addition to disparate syndromic vs non-syndromic classifications, *DIAPH1* variants are associated with differences in hearing loss frequency profiles and differences in outer hair cell function. The *ttaa* variant stands out for hearing loss that begins at low frequencies (Lalwani *et al.*, 1998), while all other variants are linked to hearing loss that begins in the high frequency range (Kang *et al.*, 2017; Neuhaus *et al.*, 2017; Kim *et al.*, 2019). This difference does not map onto the classification that we propose here, and it is unknown what might underlie high- vs low-frequency onset. Extensive electrocochleography has not been reported in all studies of *DIAPH1*-related hearing loss. Recently, a missense mutation (Glu1184AlafsTer11) in which the entirety of the DAD domain of DIAPH1 is truncated (MDxLLxxL and RRKR motifs) was linked to autosomal dominant auditory neuropathy (Wu *et al.*, 2020). The classification of auditory neuropathy was made based on preserved distortion-product otoacoustic emissions and cochlear microphonic potentials, with absent auditory brainstem response in affected individuals (Wu *et al.*, 2020). However, the Δ*ag* mutation was associated with loss of both transiently-evoked and distortion-product otoacoustic emissions (Neuhaus *et al.*, 2017), indicating a loss of outer hair cell function and a notable difference from the Glu1184AlafsTer11 variant. Therefore, while it is possible that the classification arising from this study could coincide with syndromic vs non-syndromic DFNA1 variants (see above), it does not appear that this classification correlates with differences in frequency profiles or outer hair cell function.

The number of mapped likely-pathogenic *DIAPH1* variants associated with hearing loss has increased rapidly in recent years. In addition to the five variants characterized in this paper and the microcephaly mutations described above, there are at least three other variants that are classified as ‘pathogenic’ with a hearing loss phenotype in the Deafness Variation Database (Azaiez *et al.*, 2018). The protein mutations that result from these variants are: G184A, L221F, and K847R(*fs*Ter28). To our knowledge, there are no detailed reports of the clinical features of these variants. In addition to point mutations and frameshifts, *DIAPH1* was the locus with the greatest number of exon copy number variations (CNVs) among 80 deafness genes sequenced in 80 individuals with hearing loss (Ji *et al.*, 2014). In the same study, the missense DIAPH1 variant I700N was found in one case to be accompanied by a CNV gain of exon 16 (NM_005219.5) and is annotated on the Deafness Variation Database as ‘likely pathogenic’. The mutation is located in a predicted linker between poly-pro tracks in the FH1 domain. The predicted transcript, accounting for the CNV would encode approximately 20 poly-pro tracks instead of ten in the WT protein. We would not expect this to affect autoinhibition but have not tested it directly. A variant that leads to in-frame skipping of exon 27 and therefore complete removal of the DAD domain (p.E1192_Q1220del) has been associated with more severe MTP than 1213x as well as hearing loss (Westbury *et al.*, 2018). Based on sequence, this variant is most similar to the auditory neuropathy variant p.Glu1184AlafsTer11 (Wu *et al.*, 2020). However, Wu *et al.* report no hematological findings for p.Glu1184AlafsTer11-linked auditory neuropathy (Wu *et al.*, 2020).

The native function of DIAPH1 in the inner ear is not well-understood. Particularly surprising is that, despite the wide-spread expression of DIAPH1, the auditory system and megakaryocytes are uniquely vulnerable to DIAPH1 constitutive activation, as is the case for 1213x and Δ*ag* variants. Our data show that INF2-NT does not inhibit these DIAPH1-FFC variants. It is possible that they are inhibited in trans by other formins. Mouse models have been invaluable in understanding human deafness variants, and a transgenic mouse expressing DIAPH1-1213x shows progressive hearing loss with loss of ≈10 and 40% of inner and outer hair cells, respectively, between 10- and 25-weeks (Ueyama *et al.*, 2016). Surprisingly, 8-week-old 1213x-expressing mice show greater hair cell damage following sound exposure (specifically loss of ribbon synapses and hair bundle abnormalities), but these changes were not correlated with permanent threshold shifts (Ninoyu *et al.*, 2020). To date, no mouse model has been reported for the other pathogenic DFNA1 variants.

Through advances in genome and exome sequencing, the number of *DIAPH1* variants will most likely continue to grow, and both molecular and organism-level studies will be important for understanding these forms of hearing loss. Through a systematic analysis of five variants, we report here highly disparate levels of autoinhibition, indicating that these variants could have distinct activation levels *in vivo*. Recently reported mouse models of DFNA1 will hopefully uncover the cellular mechanisms of *DIAPH1*-associated pathology in the inner ear (Ninoyu *et al.*, 2020). Thus far, those efforts have been focused on 1213X, which appears to date to be the most commonly occurring variant in the human population. The experiments reported here motivate creation of a mouse model for one of the variants in the weak or no activation class (*ttaa*, I530S, A265S). This work will be crucial for understanding DIAPH1 function in the auditory system and for treating DFNA1-associated deafness.

## Acknowledgements

The authors are grateful to Rachel Austin, Elizabeth Olson, and members of the Olson lab for helpful discussions and to Carla Hachicho for help with protein purification. The CD spectrometer was supported by NIH award 1S1OD025102-01. CLV and RL are supported by grant R15DC016462 from the NIH-NIDCD. CCC and TJ were supported by the Bernice Segal fund from the Barnard College Department of Chemistry, and AM was supported by the Sherman Fairchild Foundation.

## Accession IDs

Human DIAPH1 constructs were made from Transomic clone BC117257 and are referenced according to the numbering for NP_005210.3 (see Table S1). Human INF2 protein corresponds to NP_071934.3, and *S. pombe* profilin to NP_593827.1.

---

**Figure S1.**
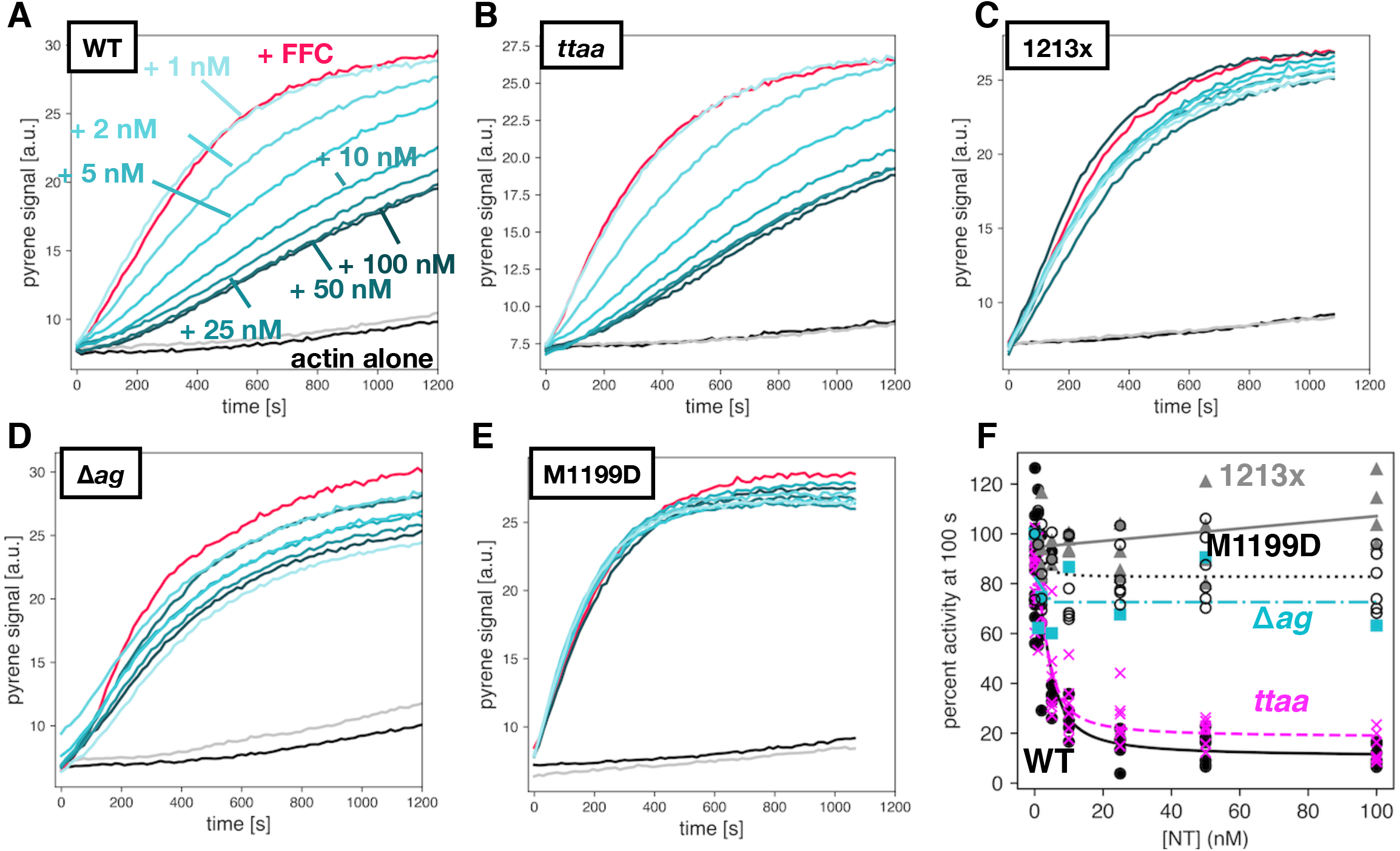
Varied auto inhibition of DIAPH1-FFC mutants by INF2-NT. Pyrene-actin assembly assays were carried out with 2 μM actin (5% pyrene-labeled), ± 5 nM of FFC, either (A) WT; (B) *ttaa*;(C) 1213x; (D) Δ*ag*; or (E) M1199D, with varied concentration of INF2-NT. Concentrations of NT are indicated in (A), and the color scheme is the same in panels (B-E), with actin plus 200 nM NT and no FFC in gray. (F) Inhibition curves calculated from raw pyrene traces and fit with a quadratic binding model that assumes 1:1 binding between dimers.

**Figure S2.**
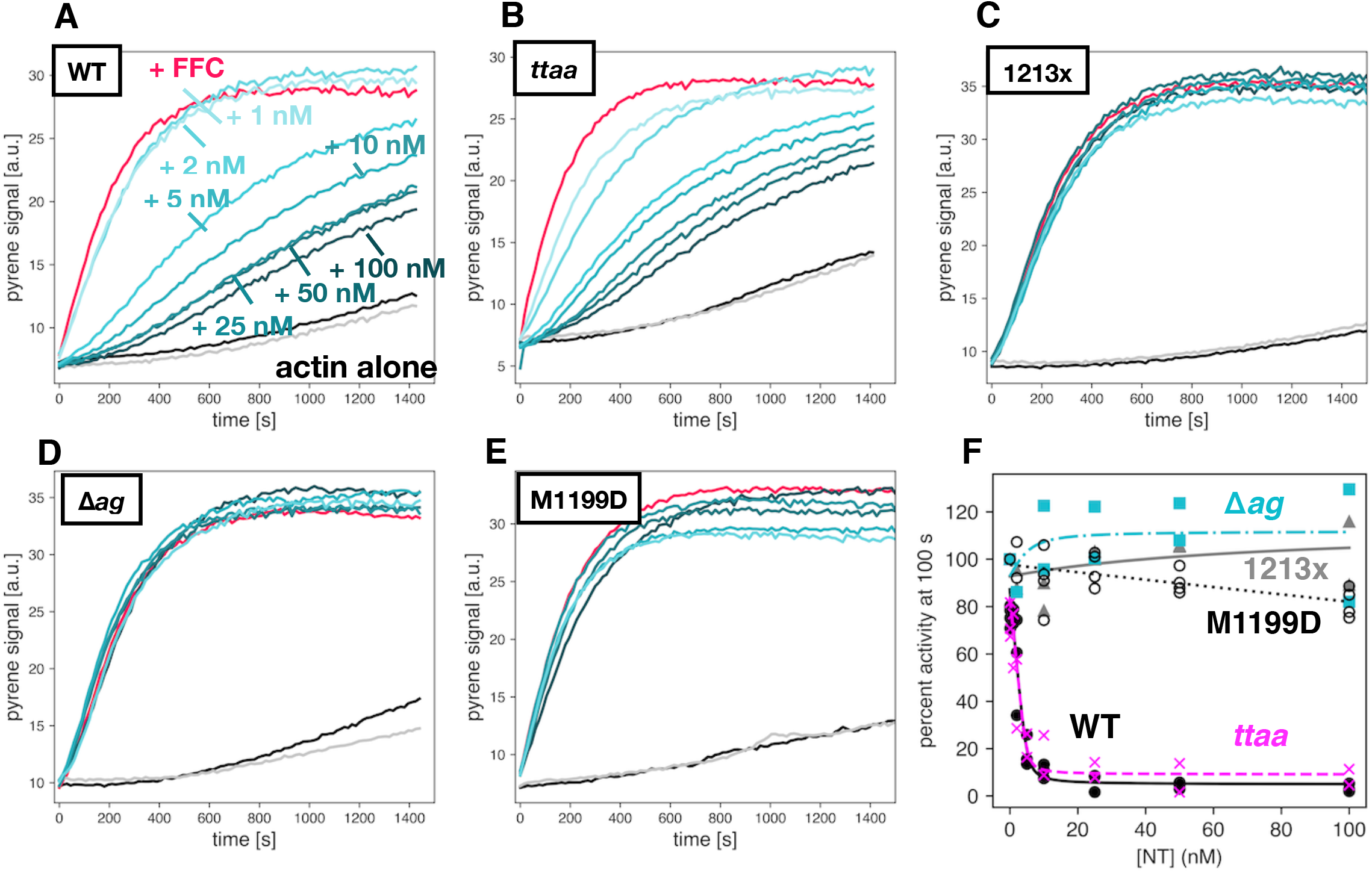
Varied auto inhibition of DIAPH1-FFC mutants. *Replicate of Figure 4 with different protein preparations.* Pyrene-actin assembly assays were carried out with 2 μM actin (5% pyrene-labeled), ± 5 nM of FFC, either (A) WT; (B) *ttaa*;(C) 1213x; (D) Δ*ag*; or (E) M1199D, with varied concentration of WT DIAPH1-NT. Concentrations of NT are indicated in (A), and the color scheme is the same in panels (B-E), with actin plus 200 nM NT and no FFC in gray. (F) Inhibition curves calculated from raw pyrene traces and fit with a quadratic binding model that assumes 1:1 binding between dimers.

**Figure S3.**
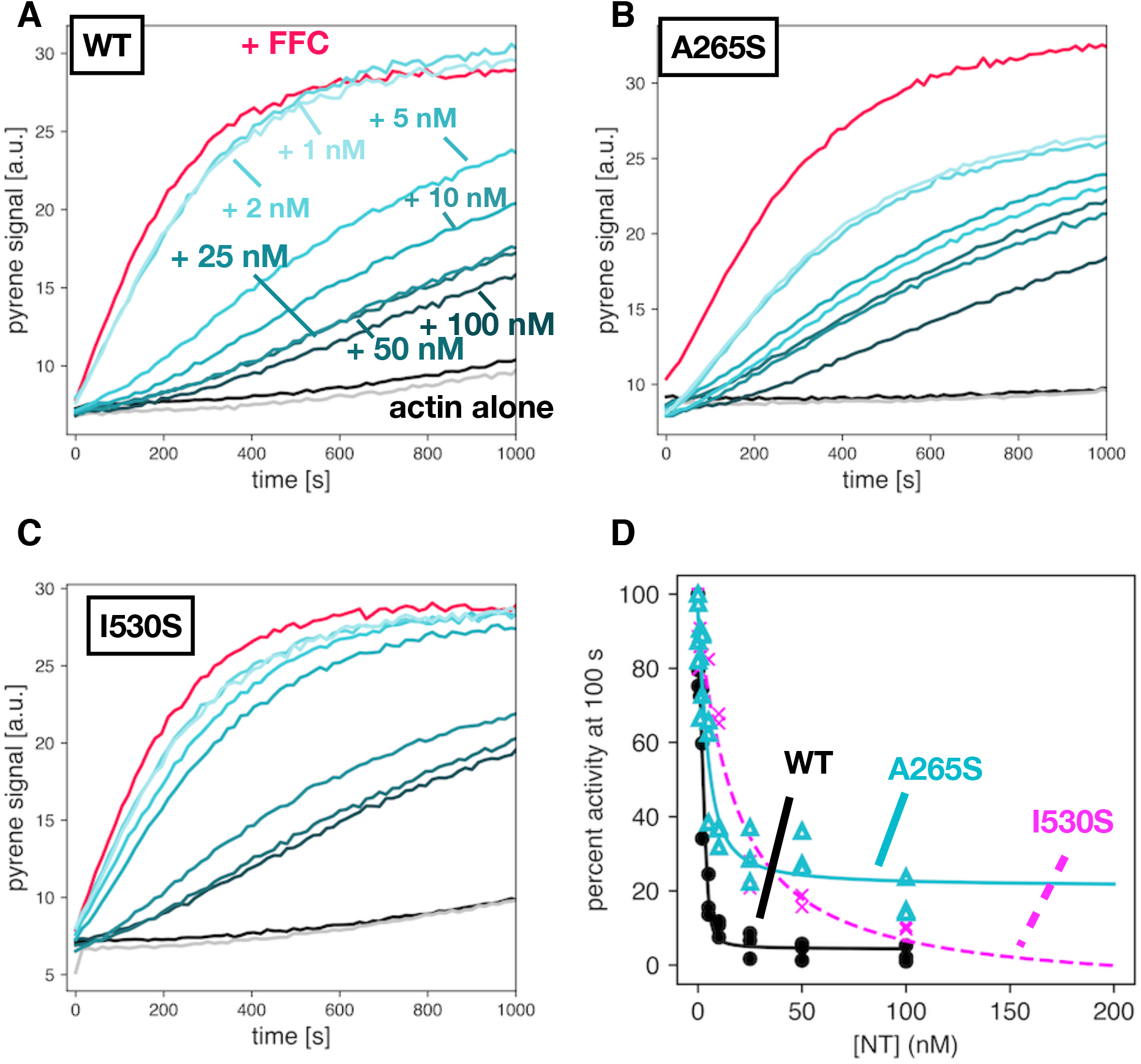
Auto inhibition by DIAPH1-NT mutants. *Replicate of data in Figure 5 with different protein preparations.* Pyrene-actin assembly assays were carried out with 2 μM actin (5% pyrene-labeled), ± 5 nM of WT-FFC, titrated with either (A) WT; (B) A265S; or (C) I530S DIAPH1-NT. Concentrations of NT are indicated in (A), and the color scheme is the same in panels (B,C), with actin plus 200 nM NT (no FFC) in gray. (D) Inhibition curves calculated from raw pyrene traces and fit with a quadratic binding model that assumes 1:1 binding between dimers.

**Table S1.**
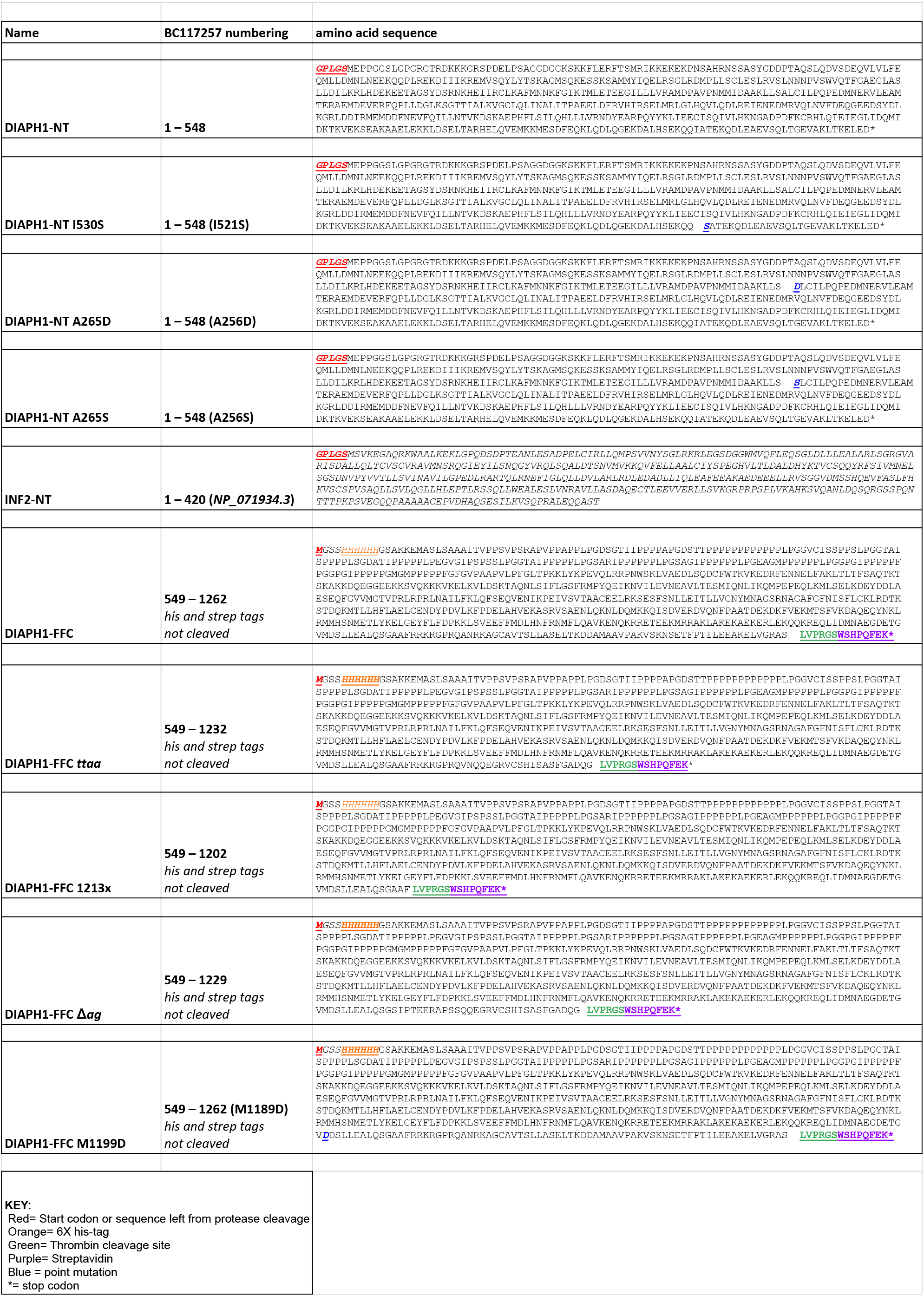
Sequences for proteins used in this study.

